# Tick Genome Assemblies: Overcoming biological limitations through advances in sequencing technologies

**DOI:** 10.1101/2025.11.11.687924

**Authors:** Katie C. Dillon, Julia C. Frederick, Hein Sprong, Travis C. Glenn, Isobel Ronai

## Abstract

Ticks are blood-feeding arthropods with approximately 1,000 species, however, only 24 species currently have a genome assembly. These genome assemblies are important resources to advance tick biology and control of tick-associated diseases. Generating tick genome assemblies is challenging due to their small body size (low DNA input), DNA contamination (from microbiota and host bloodmeals), large genome size (on average 2.4 Gbp for hard ticks), and abundant transposable elements (at least 68% of the assembly for *Ixodes* species). Advances in sequencing technologies have driven an increasing number of tick assemblies from 2011 to 2025. We characterize and assess the 54 tick genome assemblies within public genome databases using QUAST-LG and BUSCO compleasm. Then we evaluate the impact of biological source material and sequencing platforms on these tick genome assemblies. From the 54 tick assemblies, we identify 34 high-quality assemblies from 21 species that are suitable for downstream analyses. We recommend future tick genome assemblies use long-read sequencing platforms and Hi-C scaffolding to improve genomic resources for these unique blood-feeding parasites.

## THE IMPORTANCE OF TICKS

Ticks are hematophagous mites with a global health impact on humans, domesticated animals including livestock, and wildlife. As blood-feeders, ticks are vectors, transmitting a diverse range of disease agents, including viruses (for example, tick-borne *Flaviviridae*, *Dabie bandavirus*, and *Orthonairovirus*), bacteria (for example, *Borrelia* spp*., Anaplasma* spp. and Spotted Fever Group *Rickettsiae*), protozoa (for example, *Babesia* spp., *Theileria* spp.), and filarial nematodes^1,2^. The occurrence and distribution of tick-associated diseases is increasing due to urbanization and climate change^3,4^. Furthermore, ticks are developing resistance to current control strategies^5,6^.

Ticks (order Ixodida) are comprised of three extant families: hard ticks (Ixodidae), soft ticks (Argasidae), and Nuttalliellidae. While there are approximately 1,000 tick species^7,8^, few have whole-genome assemblies (DNA sequence of the autosomes and sex chromosomes; in ticks the heterogametic sex with different sex chromosomes is the male^9^) available on databases. This lack of tick genome assemblies is in part due to the unique biology of ticks.

### Tick biology considerations for genome sequencing

Key aspects of tick biology should be taken into consideration prior to whole-genome sequencing. Most notably, ticks have large genomes that are more similar in size to mammalian genomes than other vector species. We identified 17 genome assemblies of human disease vector species, spanning six taxonomic orders, from the World Health Organization’s list of major human vector-borne diseases globally (https://www.who.int/news-room/fact-sheets/detail/vector-borne-diseases). For these 17 vector assemblies, the five largest assemblies all belong to ticks (Fig. 1). These tick assemblies have an average size that is up to 19 times larger than the other vector species, such as the human body louse (Fig. 1). The large genome-size of ticks has hindered the generation of genome assemblies, as large amounts of sequencing and bioinformatic resources are required to achieve high-quality assemblies with sufficient genome coverage (the number of times a single nucleotide has been sequenced).

**Figure 1.**
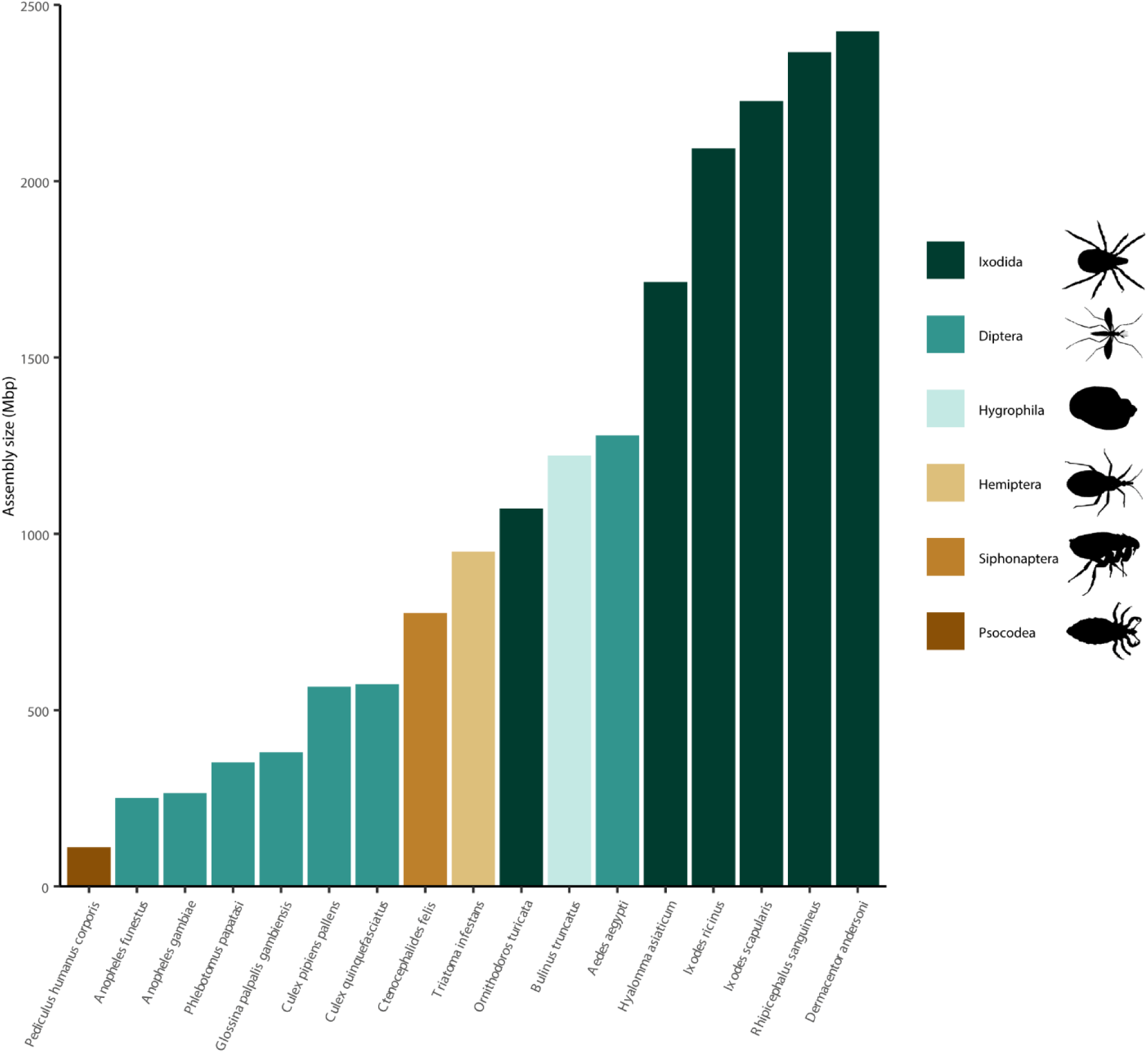
**Ticks (Ixodida) have the largest genome assembly sizes when compared with other major human disease vectors**. Genome assembly size (from the NCBI database) for 17 species, spanning six taxonomic orders, from the World Health Organization major human vector-borne diseases list (https://www.who.int/news-room/fact-sheets/detail/vector-borne-diseases). For human vector-borne diseases associated with multiple vector species, the species associated with the highest number of disease cases and has an available assembly is shown. When a vector species has multiple assemblies, the current reference assembly from NCBI is used. Species are ordered by assembly size, from smallest to largest.

Tick genomes are complex, containing numerous highly repetitive sequences. Transposable elements (mobile DNA sequences that move around the genome^10^) are nearly 70% of some tick genome assemblies, which is amongst the highest observed in animals^11^. This high proportion of transposable elements in tick genomes increases their size and complexity, complicating genome assembly.

The body size of a tick is relatively small for an animal. Adult ticks are typically only a few millimeters when unengorged, resulting in a low total DNA yield from an individual tick^12^. As tick genomes are large and highly repetitive, a relatively large amount of high molecular weight genomic DNA is required to meet DNA input minimums for genome sequencing. The first tick assemblies obtained sufficient amounts of DNA by pooling ticks^13^. In 2020, the minimum amount of high molecular weight genomic DNA required for sample preparation was achieved with single-tick DNA extractions^14^. Therefore, sequencing of individual ticks to produce high-quality tick genome assemblies has only become technically feasible within the past five years.

A DNA extraction from a tick contains contaminants due to DNA from their hosts and microbes. Typically tick species consume at least three blood meals from a host vertebrate throughout their lifecycle, potentially from multiple host species. These bloodmeals also allow the transfer of microbes from the host vertebrate to tick^15^.

Furthermore, tick species possess facultative and obligate endosymbionts^16,17^. This DNA from other organisms, as well as environmental DNA means that DNA sequencing from tick biological material generates many reads of non-tick origin (for example, 11% of total reads for an *I. ricinus* assembly^11^). In summary, generating high-quality tick genome assemblies is challenging due to their unique biology.

### Survey of tick genome assemblies and quality assessment

We conducted a survey for tick whole-genome assemblies (keyword “Ixodida”) in the publicly available sequence databases: United States National Center for Biotechnology Information (NCBI); European Nucleotide Archive (ENA); and China National Center for Bioinformation (CNCB). A total of 70 tick genome assemblies: found on NCBI (65 assemblies) or ENA (one assembly), as of April 30, 2025; found on CNCB (one assembly), as of February 28, 2025; or came from in-house (three assemblies), which were not yet publicly available. We identified 16 assemblies on NCBI that have been mistakenly classified as tick genomes; they are endosymbiont genomes or transcriptomes (Table S1). Therefore, we characterize 54 genome assemblies, representing 24 species of tick (Table S2). The tick species designations were obtained from genome assembly publications, genome databases, or personal communications.

For all 54 tick genome assemblies we assessed quality metrics for contiguity (auN, N50, L50, and L90) using QUAST-LG v.5.2.0^18^. We also assessed each assembly’s completeness with BUSCO (Benchmarking Universal Single-copy Orthologs; a BUSCO complete score is the sum of the single copy and duplicated genes that can be completely aligned to the assembly with one or more than one gene copy present) lineage arthropoda_odb10, using compleasm v.0.2.5^19^. We evaluated how the biological source material, sequencing platforms, and annotation impact tick genome assemblies.

## BIOLOGICAL SOURCE MATERIAL FOR GENOME SEQUENCING OF TICKS

We compiled the metadata of the biological material for the 54 tick genome assemblies. For the zoogeographic origin^20^ of the tick material, the most assemblies (∼35%) are generated from ticks collected in the Nearctic zoogeographic region (North America), followed by ∼31% Palearctic (Europe and China) and ∼26% Afrotropical (Africa) (Fig. 2A, Table S3). We also note that some tick species are invasive and have been collected outside their native zoogeographic region and a genome assembly generated from this material, such as *Haemaphysalis longicornis* (see ^21,22^). Currently, there are no genome assemblies from ticks collected in the Oriental region (South Asia and Southeast Asia) nor the Neotropical region (Central America and South America) (Fig. 2A, Table S3), which has the second and fourth highest number of hard tick species^20^, respectively. We recommend the Neotropical and Oriental zoogeographic regions be prioritized in future tick genome projects.

**Figure 2.**
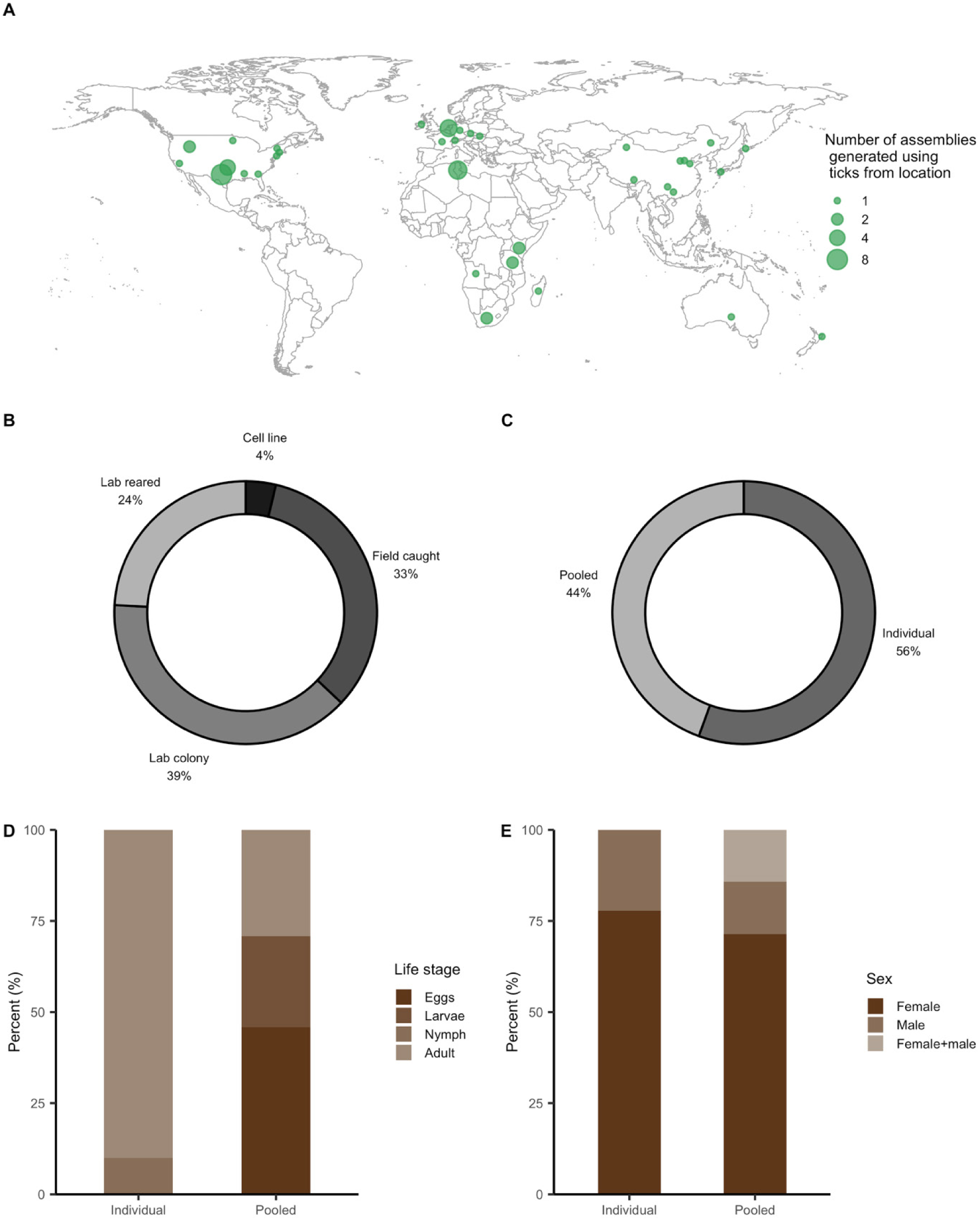
Biological source material for 54 tick genome assemblies. **A)** Geographic origin of the tick material. Two assemblies were generated using ticks from three locations. **B)** Collection source of the tick material. Laboratory colony ticks have been maintained across generations in a lab environment. We define laboratory reared ticks as those collected from the field and maintained in the laboratory until a desired life stage has been obtained, not across generations. Field caught ticks are collected from either the environment or hosts. Tick cell lines are preserved cells, typically derived from embryos. **C)** Sample size of the tick material, either an individual tick or pooled ticks. **D)** Life stage of the ticks in individual versus pooled material. **E)** Sex of the adult ticks in individual versus pooled material. See Table S3 for further details.

For the collection source of the tick material, the highest number of assemblies are generated from tick colonies (Fig. 2B, Table S3). The second highest number of assemblies are generated from ticks caught in the field and the third highest from laboratory reared ticks that were caught in the field and maintained in the laboratory until a desired life stage had been obtained (Fig. 2B, Table S3). The smallest number of assemblies are from tick cell lines (Fig. 2B, Table S3). These cell lines are typically created from heterogenous embryonic cells and maintained indefinitely in a laboratory, which leads to large-scale genome duplications, rearrangements, and deletions^23^. We recommend the biological source material for assemblies be obtained from laboratory colonies, laboratory reared, or field caught.

Tick material has varied in sample size for assemblies, with ∼56% from individual ticks and ∼44% pooled ticks (Fig 2C, Table S3). For assemblies generated from individuals, all are adult ticks, apart from three nymphs (Fig. 2D, Table S3). For assemblies generated from pooled tick material, the highest number of assemblies came from eggs (∼46%) (Fig. 2D, Table S3). For the assemblies generated from individuals, ∼78% were from adult females (Fig. 2E, Table S3), most likely because they generally are larger than males resulting in higher DNA yields. Since 2020 assemblies (contig) generated from a single individual has been feasible (Table S2-S3), often in conjunction with pooled DNA for high-throughput chromosome conformation capture (Hi-C) assembly scaffolding^24^. While pooling individual ticks or eggs provides sufficient DNA yield for a variety of sample preparation and sequencing approaches, due to variation in heterozygosity amongst individual ticks, pooling ticks leads to a less contiguous assembly^25^. We recommend prioritizing tick assemblies generated from individual ticks, to achieve a high-quality haploid assembly.

## TICK WGS AND ASSEMBLY DEVELOPMENTS FROM 2011–2025

We traced the history of the 54 tick genome assemblies. The first tick assembly (JCVI_ISG_i3_1.0) was for the species *I*. *scapularis*, with the assembly publicly available on the NCBI genome database in 2011 (Fig. 3, Table S2) and paper published in 2016^13^. This genome project resulted from a 2004 community white paper, with *I*. *scapularis* prioritized due to its medical importance in the United States, as the key tick species associated with Lyme disease^26^. The JCVI_ISG_i3_1.0 is the only tick assembly to use Sanger dideoxy chain termination on capillary instruments (first-generation sequencing)^27^. The modest sequence read lengths of Sanger was unable to produce a highly contiguous assembly due to requiring pooled individuals for input material and tick genomes being highly repetitive.

**Figure 3.**
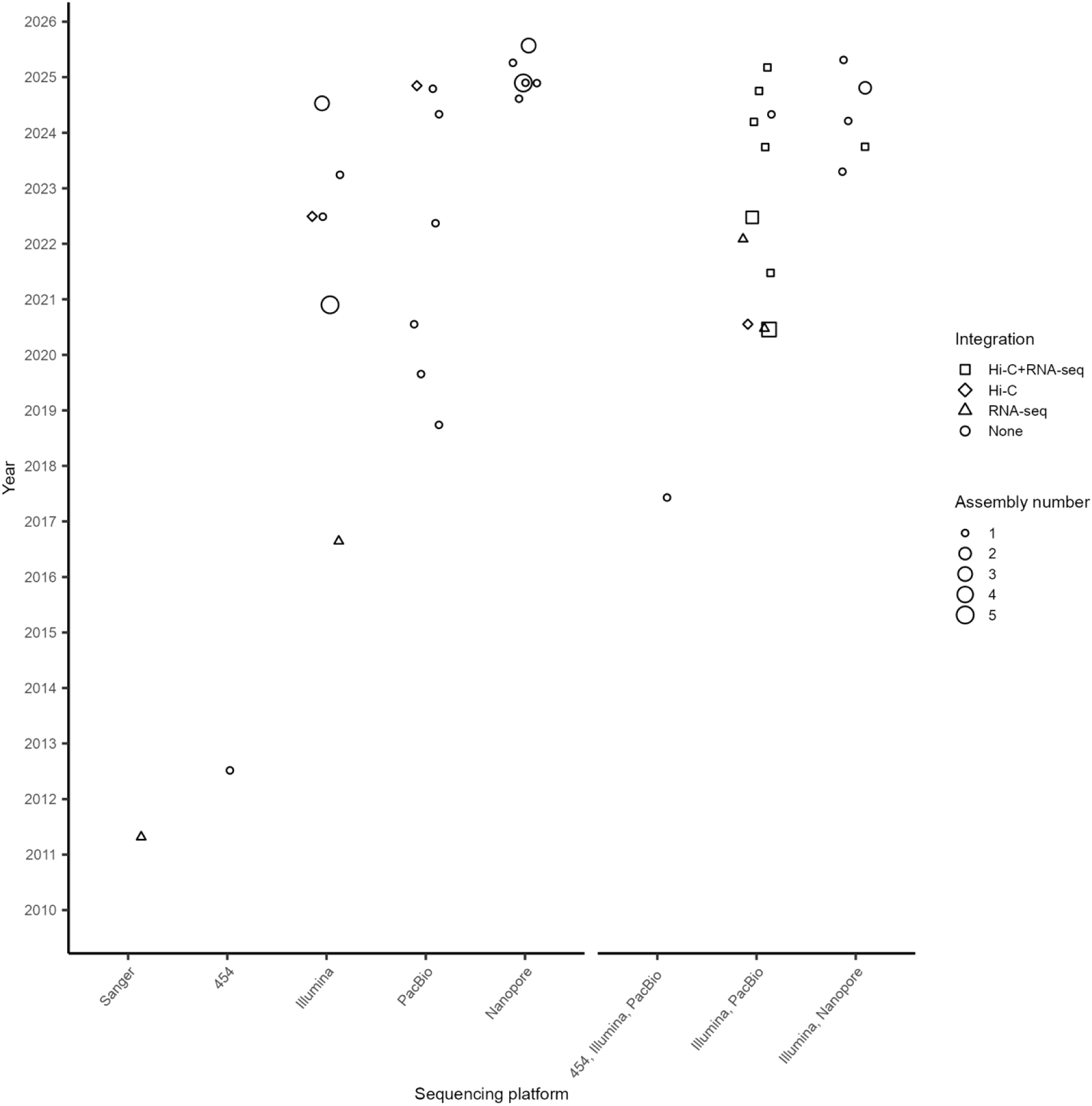
Timeline of 54 tick genome assemblies, first release was in 2011. Each point is the release date on a database of a genome project; a project has included up to five tick assemblies. Sequencing platforms are in chronological order, with first generation (Sanger or 454-FLX), second generation (Illumina) and third generation (PacBio or Nanopore). Followed by combinations of sequencing platforms. Integrations to the sequencing platforms were none, RNA-seq alone, Hi-C alone or Hi-C with RNA-seq. See Table S2 for further details.

In the 2010s tick genome assemblies started using second generation sequencing. Assemblies CCG_Rmi1.0 and Rmi2.0 of the tick *Rhipicephalus microplus* used 454-pyrosequencing (Fig. 3, Table S2). This sequencing platform obtains millions of ∼500 nt reads at a time, reducing costs compared to Sanger sequencing, but also had limitations in handling repetitive DNA^27^. In 2016, the first tick assembly using the Illumina short-read sequencing platform was released (Fig. 3, Table S2). Illumina reads are most commonly paired-end 100 or 150 nt (i.e., 200 or 300 nt total) and sequencing is massively paralleled for high throughput, providing higher genome coverage^27^. As Illumina uses short reads, it is unable to reliably map or assemble the long repetitive regions present in tick genomes.

In 2017 the first tick assembly using a third-generation sequencing platform, PacBio, was released (Fig. 3, Table S2). PacBio has a specialized library preparation (double-stranded DNA is circularized via SMRTbell adapters) and single-molecule sequencing to generate long reads (up to ∼100 kb for single-pass CLR reads or modal length of ∼20 kb when sequenced repeatedly for HiFi reads), which can handle the repetitive genomes of ticks. Initially, PacBio CLR reads were used for tick assemblies to improve contiguity^28^ and Illumina reads were often used in combination to correct uncalled or mis-called bases (Fig. 3, Table S2). The current PacBio standard is HiFi reads, which has improved base-calling accuracy and reduced the need for Illumina error correction^29^. Between 2020 to 2023, 22 tick assemblies were released, and over half used the PacBio sequencing platform (Fig. 3, Table S2).

In 2023, the first tick assembly using the Nanopore sequencing platform (another third-generation sequencer) was released (Fig. 3, Table S2). This platform generates long reads (50 bp to > 4 Mbp) in real time via flow cells with nanopores^30^. While Nanopore base-calling accuracy has greatly improved over time, ∼33% of Nanopore tick assemblies have been generated in combination with Illumina short-reads to correct base-calling errors (Fig. 3, Table S2). In 2024, 20 tick assemblies were released, the most for a single year. These assemblies were ∼40% Nanopore with the remaining using combinations of Nanopore or PacBio, with Illumina (Fig. 3, Table S2). By April 2025, six assemblies had been released, with four using Nanopore (Fig. 3). The accuracy of Nanopore reads has now improved sufficiently that Illumina reads are often no longer necessary for error correction^31^.

The development of new sequencing platform generations has been critical for overcoming the limitations of tick biology (particularly, the large genome sizes and high proportion of repeats), leading to an increasing number of tick genome assemblies.

Standalone first-generation sequencing generated ∼4% of tick assemblies, second-generation sequencing ∼20%, and third-generation sequencing ∼39% (Fig. 3, Table S2). In addition, ∼37% of tick genome assemblies have been generated using combinations of different generation sequencing (Fig. 3, Table S2). In summary, advances in sequencing technologies have driven a dramatic recent increase in tick assemblies from 2011 to 2025.

For the 54 tick assemblies, ∼37% used one or more integrations (Hi-C or RNA-sequencing) to the sequencing platform. Hi-C is a molecular approach that improves the contiguity of an assembly^27^, which is advantageous for the large and repetitive tick genomes. RNA-seq provides direct evidence of transcription, alongside the DNA coding sequences and isoforms^32^. For the 20 assemblies that used integrations, 55% had both Hi-C and RNA-sequencing, 30% RNA-sequencing alone, and 15% Hi-C alone (Fig. 3, Table S2). In mid-2020, the first tick assembly with Hi-C integrated was released, since then, ∼26% of assemblies have Hi-C integrated (Fig. 3, Table S2).

## LACK OF STANDARDIZED ANNOTATION OF TICK GENOME ASSEMBLIES

Annotation of genes and repetitive elements (particularly, transposable elements) in tick genome assemblies has not yet been standardized. These non-standardized annotations currently limit our ability to compare the genes and repetitive elements between tick assemblies, even for the same species.

Genome assemblies commonly include gene annotation. Tick assemblies have the number of protein-coding genes reported or the total number of genes reported, sometimes both numbers are reported (Table S2). We recommend future tick genome studies report both protein-coding genes and total genes.

Gene annotation has been performed for 30 tick assemblies, representing 18 species (Figure S1, Table S2). Four common annotation approaches have been used. First, a custom automated gene annotation pipeline created for each assembly, used on seven tick assemblies, with a median of ∼27,100 total genes and ∼26,400 protein-coding genes (Figure S1, Table S2). Second, taking a custom automated annotation pipeline with RNA-seq data (either pre-existing or generated alongside the genome assembly) integrated using gene modeling programs (such as AUGUSTUS and BRAKER^33^), used on 20 tick assemblies with a median of ∼28,500 total genes and ∼26,700 protein-coding genes (Figure S1, Table S2). Third, manual curation preceded by an automated pipeline that integrated RNA-seq and proteome data, used on one tick assembly, with ∼22,500 protein-coding genes (Figure S1, Table S2). Fourth, requesting NCBI to perform a RefSeq automated annotation (NCBI Eukaryotic Genome Annotation Pipeline^34^), which incorporates RNA-seq and proteome data from NCBI, used on seven tick assemblies with a median of ∼27,000 total genes and ∼26,900 protein-coding genes (Figure S1, Table S2). While four gene annotation approaches have been used and resulted in variable gene number estimates, the median number for total genes (15 assemblies) across the four approaches is between ∼27,000 to ∼28,500 and protein-coding genes (24 assemblies) is between ∼22,500 to ∼26,900. We recommend future tick genome studies integrate transcriptomes and the assembly be submitted to NCBI for annotation to achieve robust gene annotation.

As tick genomes are highly repetitive, annotation of these repetitive elements is important. Repetitive elements have been annotated in 25 tick genome assemblies, representing 15 species (Figure S2, Table S2). Commonly, repetitive elements are annotated using an automated pipeline, such as one or a combination of RepeatModeler^35^ and RepeatMasker^36^. An automated pipeline has been used on 23 tick assemblies with a median of ∼62% repetitive DNA, comprising ∼1.5% tandem repeats and ∼59% transposable elements (Figure S2A, Table S2). In studies where the transposable elements have been further classified, the median is ∼24% class I, ∼6% class II, and ∼2.4% unclassified (Figure S2B, Table S2). Manual curation of repetitive elements has been used on three *Ixodes* assemblies from one study^11^ with a median of 73% repetitive DNA, composed of ∼4.3% tandem repeats and ∼68% transposable elements (Figure S2A, Table S2). Further classification of the transposable elements gave a median ∼32% class I and ∼36% class II (Figure S2B, Table S2). We recommend future tick genome studies perform manual curation of the transposable elements until a robust automated pipeline is established.

## QUALITY ASSESSMENT OF TICK GENOME ASSEMBLIES

We comprehensively assessed the impact of sequencing platform and integration of Hi-C scaffolding on tick assembly quality, using metrics for contiguity (auN, N50, L50, and L90) and assembly completeness (BUSCO complete scores). Sanger and 454-FLX sequencing platforms were used for the first two tick assemblies ever released (Table S2), generating highly fragmented assemblies (Fig. 4A-D, Table S4). While the complete BUSCO score for the Sanger assembly was close to 90%, the 454-FLX partial assembly was close to 0% (Fig. 4A-D, Table S5).

**Figure 4.**
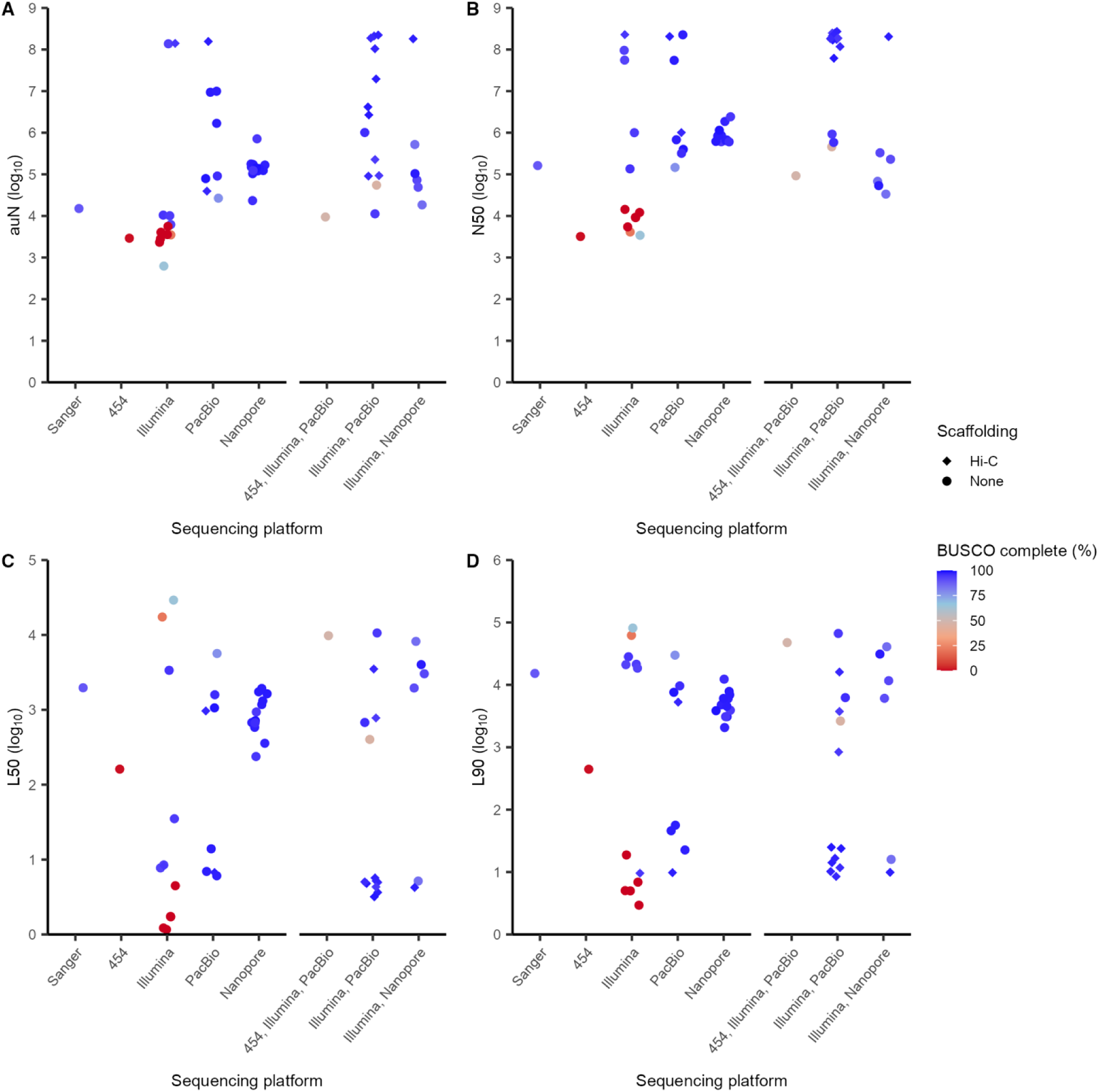
Quality metrics for 54 tick genome assemblies across sequencing platforms. **A)** auN (comprehensive metric for contiguity), as auN increases so does assembly quality. **B)** Contig N50 (the length of the shortest contig that must be included to cover half an assembly), as N50 increases, fewer longer contigs are used to cover half of the assembly, leading to a higher-quality assembly. **C)** L50 (minimal number of contigs that cover 50% of the assembly), as L50 decreases assembly quality can increase, however, the BUSCO score needs to be included in the assessment. **D)** L90 (minimal number of contigs that cover 90% of the assembly), as L90 decreases assembly quality can increase, however the BUSCO score needs to be included in the assessment. Each point is a genome assembly. Sequencing platforms are ordered by first generation (Sanger and 454-FLX), second generation (Illumina) and third generation (PacBio and Nanopore), then followed by sequencing platform combinations. Scaffolding for the sequencing platforms were none or Hi-C. BUSCO complete score (the sum of the single copy and duplicated genes that can be completely aligned to the assembly with one or more than one gene copy present), a higher BUSCO score indicates higher assembly completeness and a higher quality assembly. See Table S2, S4, and S5 for further details.

The Illumina sequencing platform has been used for 12 tick assemblies (Table S2). Only one Illumina assembly (that first constructed a 10X Genomics library) integrated Hi-C, *I. ricinus* (IXRI_v2)^37^, generating the highest quality Illumina tick assembly with a ∼95% complete BUSCO score (Fig. 4A-D, Table S4-S5). The other 11 Illumina assemblies are more fragmented and seven did not produce sufficient reads to cover the genome, resulting in lower complete BUSCO scores of 0 to ∼65% (Fig. 4A-D, Table S4-S5). Therefore, short read platforms without Hi-C are unlikely to be sufficient to handle the genome complexity of ticks.

The PacBio sequencing platform has been used for eight tick assemblies, with two integrating Hi-C (Table S2). The remaining six PacBio assemblies without Hi-C varied in quality (Fig. 4A-D, Table S4-S5). A combination of PacBio error-corrected with Illumina short-reads has been used for 13 tick assemblies (Table S2). Twelve of these assemblies have complete BUSCO scores over 92%, indicating sufficient sequencing coverage of the genome (Fig. 4A-D, Table S4-S5). Ten of these 13 assemblies integrated Hi-Cand the three without Hi-C were lower in quality regarding contiguity and coverage (Fig. 4A-D, Table S4-S5). Our findings suggest that Hi-C positively impacts assembly contiguity and quality for PacBio tick assemblies, with or without Illumina.

The Nanopore sequencing platform has been used for 12 tick assemblies (Table S2). These assemblies overlap in quality with the Illumina, PacBio, and PacBio with Illumina assemblies (Fig. 4A-D, Table S4-S5). None of the Nanopore assemblies integrated Hi-C (Fig. 4A-D). The quality and contiguity of Nanopore assemblies is moderate when compared to the other sequencing platforms. A combination of Nanopore error-corrected with Illumina short-reads has been used for six tick assemblies (Table S2).

The Nanopore with Illumina assemblies had a ∼91% average complete BUSCO value, which was unexpectedly lower than that of the standalone Nanopore assemblies with ∼97% average complete BUSCO (Fig. 4A-D, Table S4-S5). One of the Nanopore with Illumina assemblies integrated Hi-C, which had higher quality and contiguity compared to those without Hi-C (Fig. 4A-D, Table S4-S5). The quality of the remaining five Nanopore with Illumina assemblies, but without Hi-C, while similar in contiguity had lower BUSCO complete scores compared to the standalone Nanopore tick assemblies (Fig. 4A-D, Table S4-S5). Therefore, the highest quality Nanopore assembly integrated Hi-C.

Across all the sequencing platforms assemblies integrating Hi-C had higher contiguity, quality, and completeness compared to assemblies without Hi-C (Fig. 4A-D, Table S4-S5). Additionally, assemblies generated by a combination of long-read platforms (PacBio and Nanopore) then error-corrected with Illumina reads had on average, better contiguity but lower genome completeness compared to those generated by long-read platforms without error-correction, suggesting a trade-off in contiguity versus errors (Fig. 4A-D, Table S4-S5). In summary, we recommend future tick genome assembly projects use long-read sequencing platforms with integration of Hi-C scaffolding to produce high-quality assemblies.

## CHARACTERIZATION OF HIGH-QUALITY TICK ASSEMBLIES

We assessed the quality of the 54 tick genome assemblies using three metrics (number of contigs, contig N50, and BUSCO complete score) to identify 34 high-quality tick genome assemblies that are suitable for downstream analyses (Fig. 5). These high-quality assemblies have under 100,000 contigs, a contig N50 over 45 Kbp, and a complete BUSCO score over 83% (Fig. 5, Table S4-S6). The median number of contigs is ∼8,019, contig N50 is ∼1.7 Mbp, and complete BUSCO value is 96%. High-quality assemblies of other arthropod taxa have N50s over 10 Kbp and >90% complete BUSCOs^38^. The 34 high-quality tick assemblies have all been generated since 2020 (Fig. 3); the sequencing platforms are ∼15% Illumina, ∼18% Nanopore, ∼18% Nanopore+Illumina, ∼21% PacBio, and ∼29% PacBio+Illumina (Fig. 5, Table S4-S6). Approximately 41% of the high-quality tick assemblies integrated Hi-C (Fig. 5, Table S4-S6).

**Figure 5.**
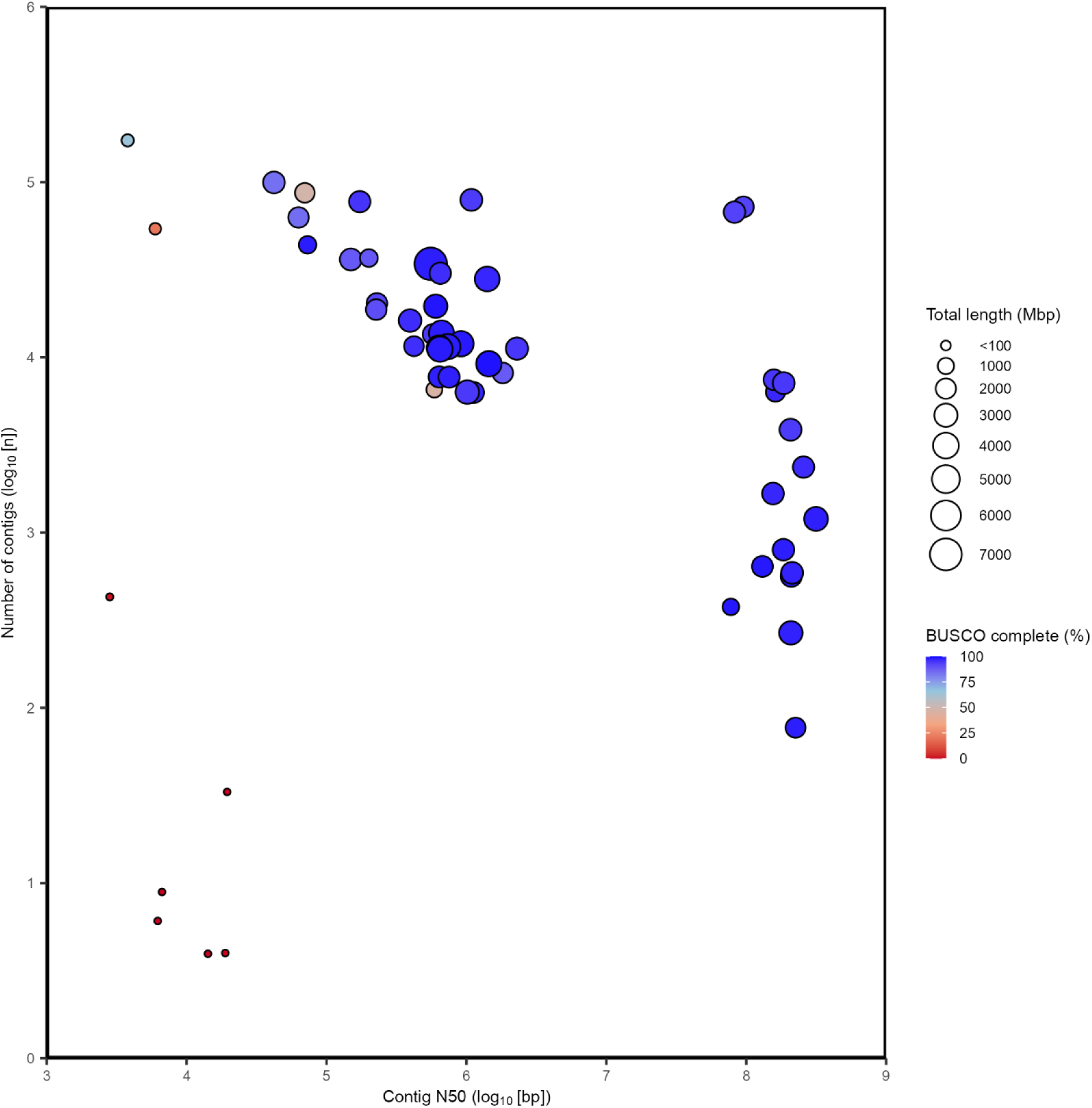
Quality assessment of 54 tick genome assemblies. Identified 34 high-quality assemblies (lower contig number, higher N50, and higher BUSCO score), including a cluster of 14 highest quality assemblies. Each dot is a genome assembly. Contig N50 is the length of the shortest contig that must be included to cover half an assembly, a higher N50 means less small contigs which is a more complete, less fragmented assembly. Number of contigs, fewer contigs typically results in a more contiguous assembly as there is less fragmentation. However, few contigs associated with a low N50 and small assembly length can lead to poor assembly quality. BUSCO complete score, a higher BUSCO score indicates higher assembly completeness and a higher quality assembly. As not all assemblies were obtained from NCBI, we used a size metric that could be calculated for all 54 assemblies: the total length of the assembly (generated by QUAST) which includes uncalled bases. QUAST total length more closely represents the haploid assembly size, not genome size. See Table S4-S6 for further details.

We also identify a cluster of 14 tick assemblies that are the highest quality, which have under 7,500 contigs, a contig N50 over 84 Mbp, and a complete BUSCO score over 93% (Fig. 5, Table S4-S6). These 14 highest-quality assemblies have all been generated since 2020 (Fig. 3); the sequencing platforms are ∼7% Illumina, ∼7% Nanopore+Illumina, ∼21% PacBio, and ∼64% PacBio+Illumina (Fig. 5, Table S4-S6). All 14 highest quality tick assemblies integrated Hi-C, except for the USDA_Rmic assembly (Fig. 5, Table S2). Our use of comprehensive metrics for the high-quality tick assemblies means future tick assembly projects now have metrics to benchmark against.

## TICK SPECIES WITH HIGH-QUALITY GENOME ASSEMBLIES

The 34 high-quality genome assemblies represent 21 species of tick (Fig. 6). As some species have multiple high-quality assemblies, we determine the best quality assembly for each of the 21 tick species (Fig. 6, Table S6).

**Figure 6.**
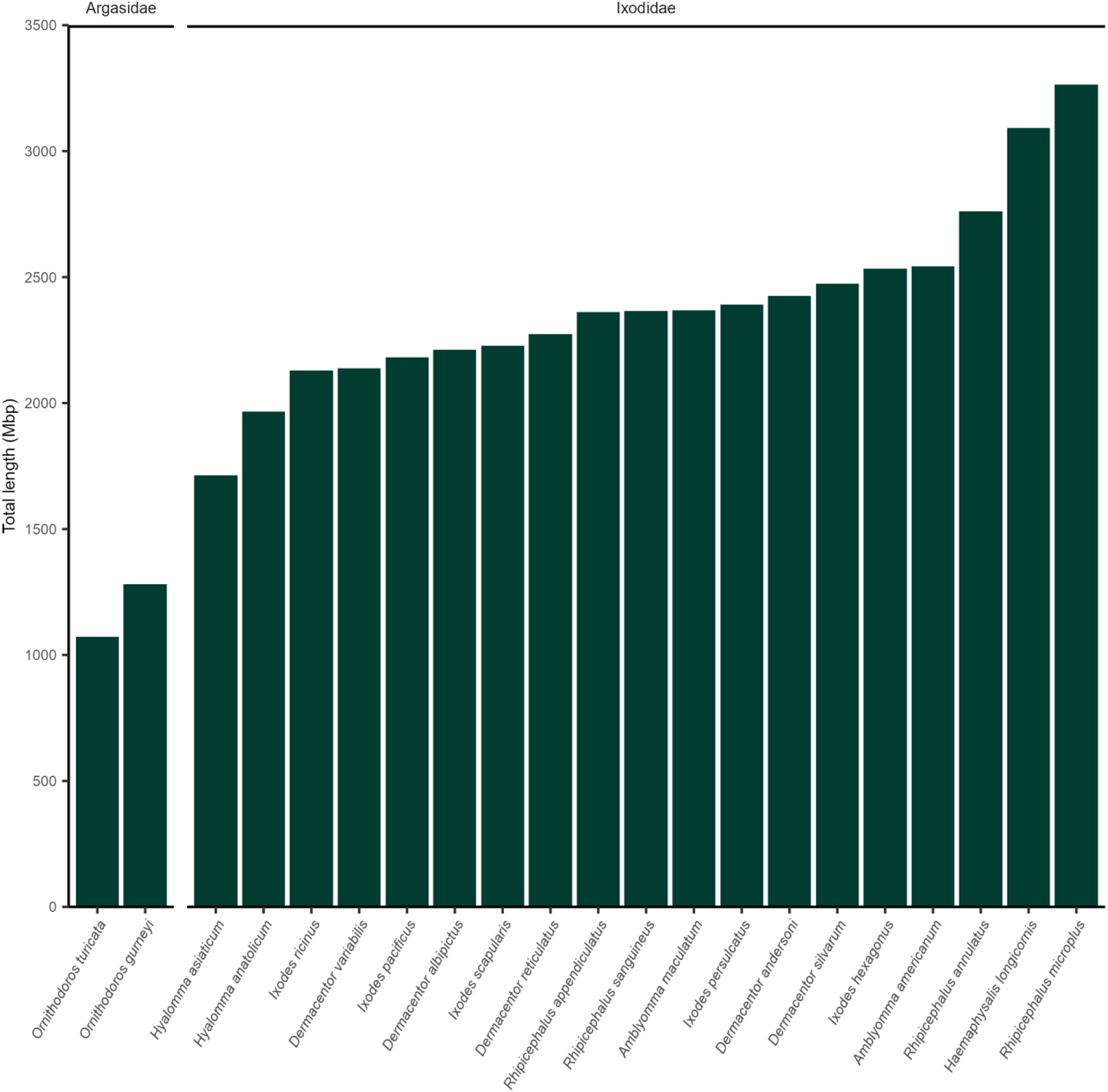
Assembly total length of the best quality genome assembly for 21 tick species. The best assembly for each tick species was identified from the high and highest quality assemblies. Tick species are ordered by assembly total length, from smallest to largest. See Table S6 for further details.

For the 21 best quality assemblies, we find hard ticks have an average assembly length of ∼2,390 Mbp and soft ticks ∼1,176 Mbp (Fig. 6, Table S4). *Rhipicephalus* spp. have the largest assembly length across all the tick species, with an average of ∼2,689 Mbp. The two *Hyalomma* spp. have the smallest assembly lengths for the hard ticks, with an average of ∼1,840 Mbp (Fig. 6, Table S4). The two soft tick *Ornithodoros* spp. have the smallest assembly lengths for all ticks, with an average of ∼1,176 Mbp (Fig. 6, Table S4). These reported assembly sizes (length of gapped DNA sequence), however, are not directly equivalent to the ‘genome size’. The average genome size of seven hard tick species is 2,671 Mbp and two soft tick species is 1,281 Mbp using flow cytometry^39^. We find tick assembly size (NCBI assembly size or QUAST assembly total length) is on average 95.5% of the genome size (Table S7), n = 16. Overall, we hypothesize that hard ticks have larger genomes compared to soft ticks (Fig. 6) due to a higher proportion of repetitive elements (Fig. S2A).

The 21 species with high-quality assemblies are two soft ticks (Argasidae) and 19 hard ticks (Ixodidae), with five *Ixodes* spp. and 14 non-*Ixodes* spp. (Table S7). These assemblies are 0.9% (2/216) of soft tick species and 2.4% (19/786) of hard tick species^7,8^. We recommend that soft tick species be prioritized for genome sequencing. So far, the tick species sequenced are those of medical and veterinary importance. Therefore, we also recommend future tick genome projects prioritize capturing the diversity of tick species.

## CONCLUDING REMARKS

We have assessed 54 genome assemblies of ticks, identifying 34 high-quality assemblies for 21 tick species. Our assessment of the tick genome assemblies has provided an opportunity to make evidence-based recommendations for future tick genome assembly projects. We recommend future assemblies better represent the biodiversity of tick species, particularly the soft ticks and from unrepresented zoogeographic regions. In addition, to obtain the highest quality tick genome assemblies, future studies should prioritize obtaining individual, adult ticks for DNA extractions and use long-read sequencing platforms integrated with Hi-C scaffolding and transcriptomes. To achieve the best quality gene and repetitive element annotations future studies should prioritize manual curation. These recommendations will hopefully be helpful for future tick genome assembly studies.

Major developments in sequencing technologies have helped overcome the biological challenges of tick genomes. However, there are genomic resources that are well established in other non-model arthropods, which are still needed for ticks. These genomic resources include: a chromosome-anchored reference genome using fluorescent *in situ* hybridization (FISH)^40^, phylogeographic representation from sequencing multiple individuals per species^41^, and cross-species whole genome alignments^42^. New sequencing technologies to further advance tick genomes include Assay for Transposase-Accessible Chromatin with sequencing (ATAC-seq)^43^ to characterize regulatory elements, and Telomere-to-Telomere (T2T) gapless sequencing^44^ to characterize structural elements. These additional approaches, alongside advancements in genome assembly software, will greatly improve our understanding of tick genomes and tick biology.

With the availability of high-quality tick genome assemblies, there are opportunities to improve our understanding of tick biology. For example, comparative genomic methods of the animal ancestral linkage groups suggest tick genomes are amongst the least derived arthropod genomes^45^. Also, the tick genome assemblies have been essential for advancing tick research, from tick vectorial capacity^46^ to anti-tick vaccines^47^. The tick genome assemblies have so far been referred to or used in over 130 follow up studies (Table S8), since 2011. These follow up studies have four key themes: identifying gene targets for anti-tick vaccination studies, tick control, and acaracide resistance; annotation of repetitive elements; intra-species and inter-species comparative genomics studies; ‘omic studies of transcriptomics and proteomics. In summary, the tick genome assemblies have enabled major advances in our understanding of tick biology. We look forward to new tick genome assemblies providing further insights into the fascinating biology of these important blood-feeding parasites.

### Ethics approval

None.

### Availability of data and materials

All supporting datasets for this study are available in the supplementary material.

## Acknowledgements

We thank all the researchers who have pushed the boundaries of tick genomics and shared with us unpublished information. In particular, we thank the following researchers for their willingness and transparency to provide information on unpublished tick genome assemblies that they have released on databases, in alphabetical order: Joshua Benoit, Seemay Chou, Christopher Faulk, Jan Forth, Melissa Klein, Michel Labuschagne, Susan Noh, Pia Olafson, Perot Saelao, Ryan O. M. Rego, Claude Rispe, Hailey Schau, and Pavlina Vechtova. We also acknowledge Roslyn Hickson for facilitating the resources that made the *O*. *gurneyi* genome assembly possible. The use of the unpublished genome assemblies in this paper does not constitute their announcement, as details related to preparation, assembly and gene annotation processes have yet to be described.

## Author contributions

Conceptualization: KCD, JCF, TCG, IR Validation: KCD

Formal analysis: KCD Investigation: KCD, JCF Resources: HS

Data Curation: KCD

Writing - Original Draft: KCD, IR

Writing - Review & Editing: KCD, JCF, HS, TCG, IR Visualization: KCD, IR

Supervision: TCG, IR

Project administration: TCG, IR Funding acquisition: HS, TCG, IR

