## Supplementary material for "Tick Genome Assemblies: Overcoming biological limitations through advances in sequencing technologies": Figure S

**Table S1. Sequence data excluded from the tick genome assemblies dataset.** We identified 16 ‘tick genome assemblies’ (Ixodida, taxon 6935) in the NCBI database as non-Ixodida based on their publications. For these non-Ixodida assemblies we provide their NCBI assembly ID, associated tick family and species, true sequence type (such as endosymbiont DNA or mRNA) and publication reference.

**Table S2. Tick genome assemblies dataset.** Tick assemblies in the National Center for Biotechnology Information (taxon: 6935), China National Center for Bioinformation, and European Nucleotide databases. We identified 54 assemblies for 21 tick species. For each tick assembly a simplified unique ID was created. We provide the tick family and species designation according to the genome publication, NCBI, or personal communication. Also, we provide the database-assigned assembly ID, source database (for NCBI assemblies the GenBank reference ID and whether a flag is present), BioProject ID, and the database release date. We classified each assembly as pseudo-haploid, as they contained high duplicates based on compleasm BUSCO duplicate score. Then we classified each assembly’s repeats as soft-masked or unmasked, based on visual inspection of the assembly and confirmation with assembly publisher. The sequencing platform used to generate sequence reads for each assembly was obtained from NCBI and the genome publication. We report whether an assembly was generated through short-reads, long-reads, and had Hi-C integration. We also report if transcriptome data was integrated. Whether assemblies have suppressed RefSeq data is reported. For each assembly the number of total genes and protein-coding genes (in square brackets) reported in the genome publication or NCBI; which were generated by a custom automated pipeline with or without RNA-seq data, manual curation combined with automated pipeline or the NCBI Eukaryotic Genome Annotation Pipeline. Repeat and transposable element content as a proportion of each genome obtained from the genome publication. The estimated genome size for each assembly obtained from the genome publication. For genome assemblies available on NCBI, the following data obtained from NCBI is reported: assembly size (Mbp), chromosome number, scaffolds, scaffold N50, contigs, contig N50, and genome coverage. The genome publication with publication date for each assembly is included, otherwise the contact person for the assembly provided.

**Table S3. Biological source material metadata for 54 tick genome assemblies.** For the genome assemblies (Table S2), the biological source metadata was determined using database information, genome publication, and personal communications with the contact person for the assembly submission. Each tick assembly has a unique ID and database-assigned assembly ID. The input material for DNA extraction used to generate the primary contig assembly was identified: the feeding status (engorged or unengorged) of the tick prior to DNA extraction; the life stage; if an adult, its sex; sample tissue; and the sample size of the tick material (individual or pooled). The region and continent that the tick(s) was originally collected and the collection source of the ticks (such as laboratory colony or field caught). If the tick assembly represents tick(s) collected from a laboratory colony, then the colony or strain ID and number of generations is provided.

**Table S4. QUAST assessment for 54 tick genome assemblies.** For the tick genome assemblies (Table S2) to assess assembly quality metrics we used QUAST-LG v.5.2.0. Each tick assembly has a unique ID and database-assigned assembly ID. The number of contigs and the total length of each assembly is reported (>= 0 bp, >= 1,000 bp, >= 5,000 bp, >= 10,000 bp, >= 25,000 bp, >= 50,000 bp). The total number of contigs (with QUAST-LG, contigs smaller than 3,000 bp are removed). The length of the largest contig is provided. Total length is the total number of bases, including Ns. GC% represents the proportion of G and C nucleotides in the assembly relative to its total length. N50 and N90 are the lengths in which all contigs equal to or longer than that length account for at least 50% or 90% of the total assembly, respectively. auN (area under the N curve), is a more comprehensive statistic than N50, as it is less affected by contig length and considers the entire Nx curve. L50 and L90 represent the smallest number of contigs that cover 50% or 90% of the total assembly, respectively. The number of Ns per 100 kbp represents uncalled and unmasked bases in the assembly.

**Table S5. BUSCO assessment for 54 tick genome assemblies.** For the tick genome assemblies (Table S2) to assess assembly completeness metrics with BUSCO we used compleasm v.0.2.5. Each tick assembly has a unique ID and database-assigned assembly ID. Fragmented genes of subclass 1 and subclass 2 can be partially aligned to the assembly where one portion cannot or can be aligned at another position, respectively. Missing genes are unable to be aligned to the assembly. A total of 1,013 BUSCO genes from the phylum Arthropoda (arthropoda_odb10) were used to assess each assembly.

**Table S6. We identify 34 assemblies as high or highest quality from the 54 tick genome assemblies.** We used the QUAST (Table S4) and BUSCO (Table S5) assessments of the 54 tick genome assemblies (Table S2) to identify 34 high-quality tick genome assemblies, including a cluster of the 14 highest quality assemblies. Each tick assembly has a unique ID and database-assigned assembly ID. The database release date for each assembly, sequencing platform, Hi-C integration is summarized from Table S2. The total number of contigs (with QUAST-LG, contigs smaller than 3,000 bp are removed), contig N50 (see Table S4), and the complete BUSCO score (Table S5). We include an explanation for our classification of an assembly as high or highest quality. Highest quality assemblies have a low number of contigs, high contig N50, high complete BUSCO score, and low to average duplicate BUSCO score. High-quality assemblies have low to average number of contigs, low to average contig N50, average to high complete BUSCO score, and low to average duplicate BUSCO score. We also identify the best assembly for each tick species.

**Table S7. Tick biology for the 21 species with high-quality genome assemblies.** For the 21 tick assemblies identified as the best quality for each tick species (Table S6), the taxonomic family and species are reported. The diploid (2n) chromosome number for both male and female ticks, alongside the sex chromosome system obtained from the scientific literature. Genome size (Mbp) has been estimated using k-mer frequency or flow cytometry obtained from the scientific literature. NCBI assembly size of the best quality tick genome assembly is given, this is calculated based on the expected genome size range of the species (https://www.ncbi.nlm.nih.gov/genbank/genome-size-check/). The QUAST total length of the highest quality tick genome assembly (Table S4), this is the total number of bases in the assembly. We calculated the ratio of the NCBI tick genome assembly size to the estimated genome size, and we also calculated the ratio of the QUAST total length of the highest quality tick genome assembly for the species to the estimated genome size of the tick species. References are included for each species' chromosomes, sex chromosomes, and genome size estimation.

**Table S8. Scientific impact of the 54 tick genome assemblies.** We searched the databases Google Scholar and Web of Science for publications that cited or referred to the 54 tick genome assemblies (Table S2). Search terms included the assembly's NCBI GenBank and RefSeq IDs, assembly ID, BioProject ID, BioSample ID, and Sequence Read Archive ID. For each publication we provide the title and DOI.

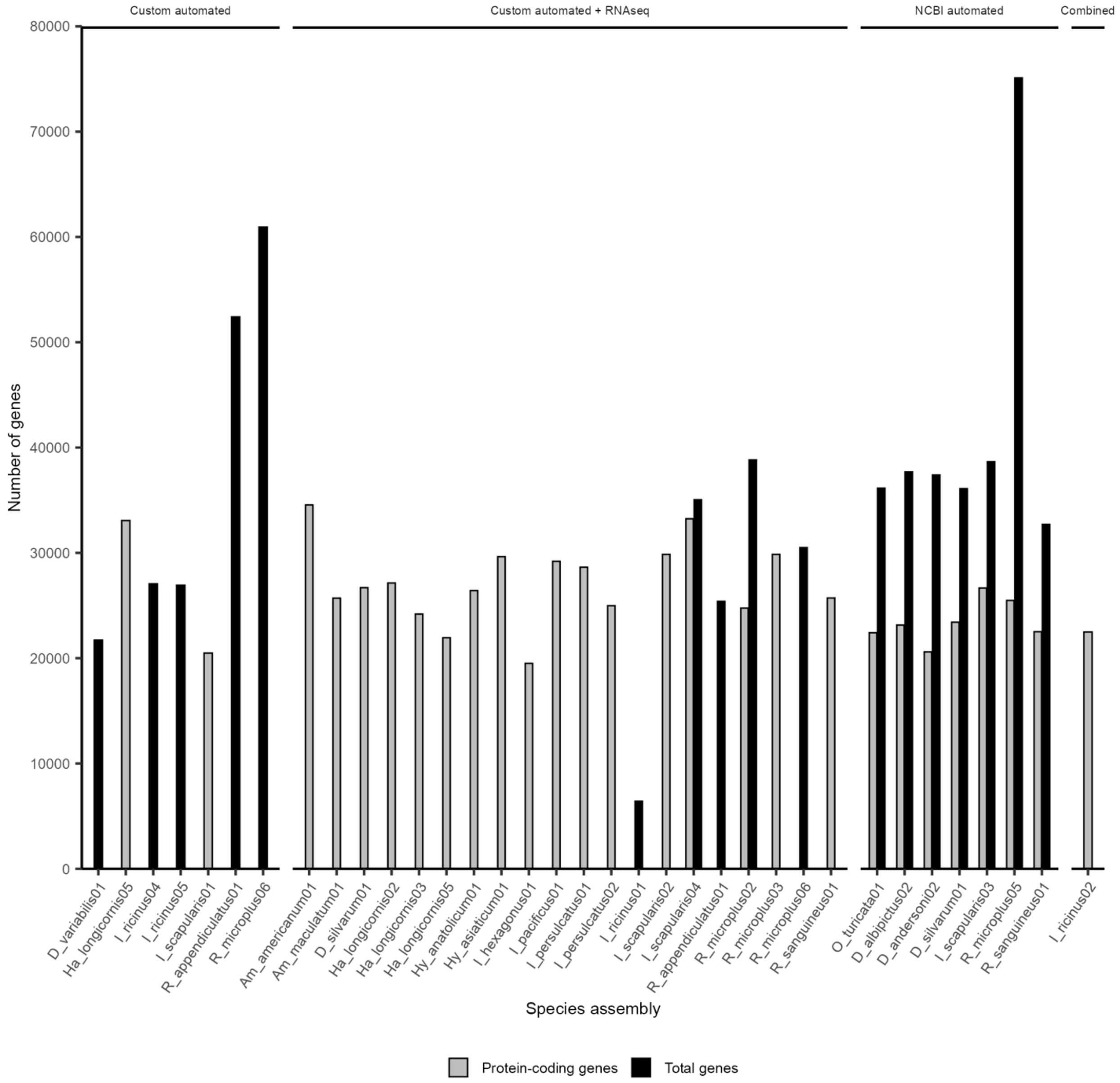


**Figure S1. Number of genes reported for the tick genome assemblies using four annotation approaches.** Thirty tick assemblies report the number of genes in the NCBI database or genome publication. Genes have been annotated with a custom automated pipeline, custom automated pipeline with RNA-seq data, manual curation combined with automated pipeline (RNA-seq and proteome data integrated), and the NCBI Eukaryotic Genome Annotation Pipeline (RNA-seq and proteome data integrated). The number of genes reported was the number of protein-coding genes and total number of genes. See Table S2 for further details.

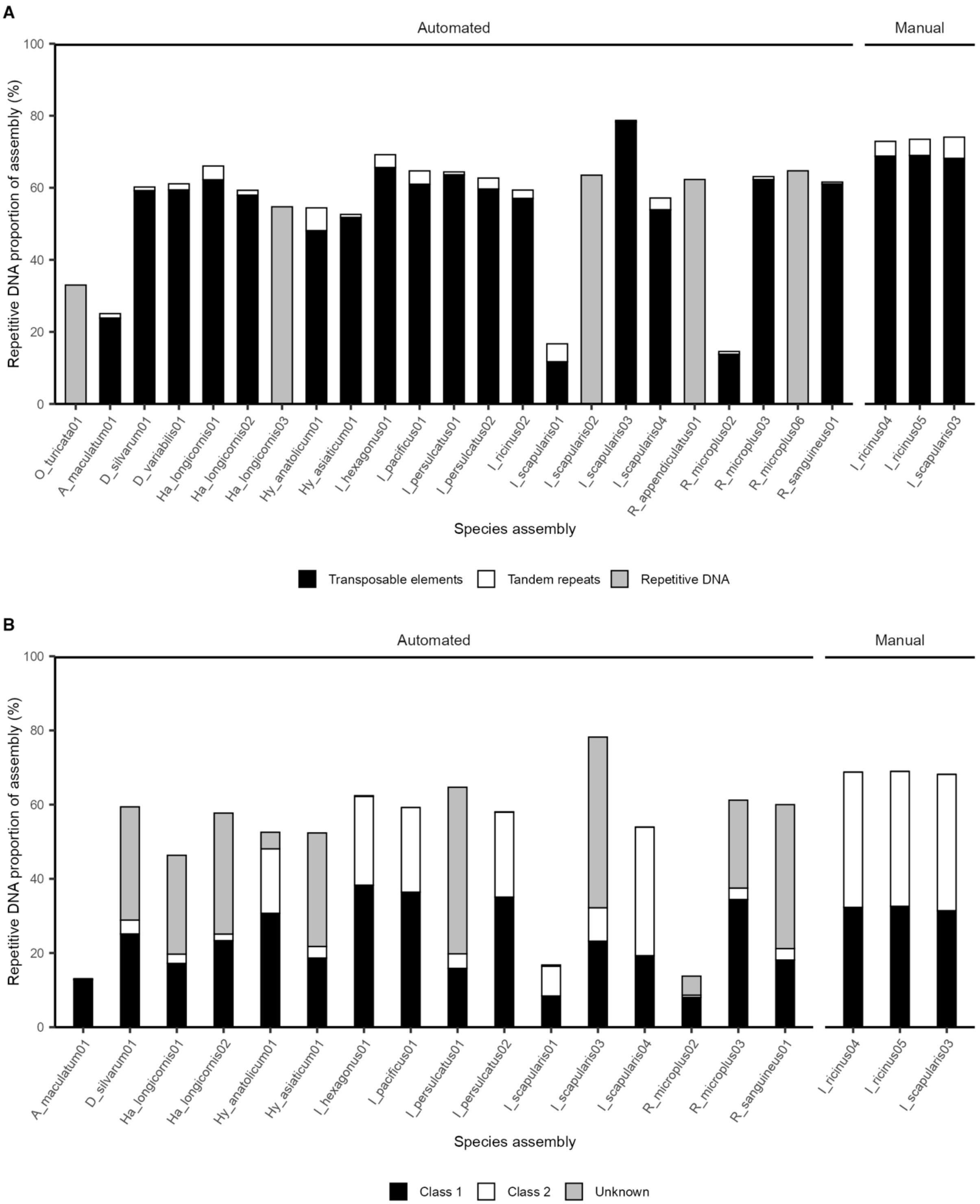


**Figure S2. Proportion of repetitive DNA (particularly transposable elements) reported for the tick genome assemblies using two annotation approaches. A)** Twenty-five tick assemblies report the proportion of repetitive elements in the genome publication. The proportion of repetitive DNA that is transposable elements or tandem repeats. Repetitive DNA has either been annotated using an automated pipeline or manually. **B)** Eighteen tick assemblies report the proportion of transposable element classes in the genome publication. Transposable elements are classified as class I, class II, or unknown. The transposable elements have either been annotated using an automated pipeline or manually. See Table S2 for further details.

**File S1. References for the 54 tick genome assemblies assessed**. Nine assemblies have publications that are in preparation and one assembly no publication is planned.

| | |
|---|---|
| O_gurneyi01 | Paper in preparation (contact Klein, M.). |
| O_moubata01 | Vechtova, P., Fussy, Z., Sterba, J., Filatov, S., Rohackova, H., Rego, R.O.M. (in prep.) Genome Sequence of the soft tick *Ornithodoros moubata* Murray 1877 (Acari: Ixodida: Argasidae): Vector of the African Swine Fever Virus and the Relapsing Fever Pathogen, *Borrelia duttonii*. |
| O_moubata02 | Forth, J.H. *et al.* (2020) Identification of African swine fever virus-like elements in the soft tick genome provides insights into the virus' evolution. *BMC biology* 18, 136. https://doi.org/10.1186/s12915-020-00865-6 |
| O_moubata03 | Forth, J.H. *et al.* (2020) Identification of African swine fever virus-like elements in the soft tick genome provides insights into the virus' evolution. *BMC biology* 18, 136. https://doi.org/10.1186/s12915-020-00865-6 |
| O_porcinus01 | Forth, J.H. *et al.* (2020) Identification of African swine fever virus-like elements in the soft tick genome provides insights into the virus' evolution. *BMC biology* 18, 136. https://doi.org/10.1186/s12915-020-00865-6 |
| O_porcinus02 | Forth, J.H. *et al.* (2020) Identification of African swine fever virus-like elements in the soft tick genome provides insights into the virus' evolution. *BMC biology* 18, 136. https://doi.org/10.1186/s12915-020-00865-6 |
| O_porcinus03 | Forth, J.H. *et al.* (2020) Identification of African swine fever virus-like elements in the soft tick genome provides insights into the virus' evolution. *BMC biology* 18, 136. https://doi.org/10.1186/s12915-020-00865-6 |
| O_turicata01 | Tietjen, M. *et al.* (2025) Genome report: whole-genome assembly of the relapsing fever tick *Ornithodoros turicata* Dugès (Acari: Argasidae). *G3 Genes\|Genomes\|Genetics* 15. https://doi.org/10.1093/g3journal/jkaf103 |
| Am_americanum01 | Chou, S. *et al.* De novo assembly of a long-read *Amblyomma americanum* tick genome. *Arcadia Science* (2023). https://doi.org/https://dx.doi.org/10.57844/arcadia-9b6j-q683 |
| Am_maculatum01 | Ribeiro, J.M. *et al.* (2023) A draft of the genome of the Gulf Coast tick, *Amblyomma maculatum*. *Ticks and tick-borne diseases* 14, 102090. https://doi.org/10.1016/j.ttbdis.2022.102090 |
| D_albipictus01 | Paper in preparation (contact Olafson, P.). |
| D_albipictus02 | Paper in preparation (contact Olafson, P.). |
| D_andersoni01 | Paper in preparation (contact Benoit, J.). |
| D_andersoni02 | Paper in preparation (contact Benoit, J.). |
| D_reticulatus01 | Dillon, K.C., Sprong, H., Baede, V.O., de Paula Baptista, R., Ray, D.A., Glenn, T.C. (in prep.) Annotated Nanopore genome |

| | |
|---|---|
| | assemblies from the tick *Dermacentor reticulatus* illuminates intra- and inter-species differences in repetitive elements. |
| D_reticulatus02 | Dillon, K.C., Sprong, H., Baede, V.O., de Paula Baptista, R., Ray, D.A., Glenn, T.C. (in prep.) Annotated Nanopore genome assemblies from the tick *Dermacentor reticulatus* illuminates intra- and inter-species differences in repetitive elements. |
| D_reticulatus03 | Dillon, K.C., Sprong, H., Baede, V.O., de Paula Baptista, R., Ray, D.A., Glenn, T.C. (in prep.) Annotated Nanopore genome assemblies from the tick *Dermacentor reticulatus* illuminates intra- and inter-species differences in repetitive elements. |
| D_silvarum01 | Jia, N. *et al.* (2020) Large-scale comparative analyses of tick genomes elucidate their genetic diversity and vector capacities. *Cell* 182, 1328–1340. e1313. https://doi.org/10.1016/j.cell.2020.07.023 |
| D_variabilis01 | Cassens, J. *et al.* (2025) The Genome of the American Dog Tick (*Dermacentor variabilis*). *G3: Genes, Genomes, Genetics* 15. https://doi.org/10.1093/g3journal/jkaf130 |
| Ha_longicornis01 | Guerrero, F.D. *et al.* (2019) The Pacific Biosciences de novo assembled genome dataset from a parthenogenetic New Zealand wild population of the longhorned tick, *Haemaphysalis longicornis* Neumann, 1901. *Data in Brief* 27, 104602. https://doi.org/10.1016/j.dib.2019.104602 |
| Ha_longicornis02 | Jia, N. *et al.* (2020) Large-scale comparative analyses of tick genomes elucidate their genetic diversity and vector capacities. *Cell* 182, 1328–1340. e1313. https://doi.org/10.1016/j.cell.2020.07.023 |
| Ha_longicornis03 | Yu, Z. *et al.* (2022) The new *Haemaphysalis longicornis* genome provides insights into its requisite biological traits. *Genomics* 114, 110317. https://doi.org/10.1016/j.ygeno.2022.110317 |
| Ha_longicornis04 | Umemiya-Shirafuji, R. *et al.* (2023) Draft genome sequence data of *Haemaphysalis longicornis* Oita strain. *Data in Brief* 49, 109352. https://doi.org/10.1016/j.dib.2023.109352 |
| Ha_longicornis05 | Moustafa, M.A.M. *et al.* (2025) Genome of the invasive North American *Haemaphysalis longicornis* tick as a template for bovine anti-tick vaccine discovery. *BMC Genomics* 26, 307. https://doi.org/10.1186/s12864-025-11477-1 |
| Hy_anatolicum01 | Wang, J. *et al.* (2024) Insight into *Hyalomma anatolicum* biology by comparative genomics analyses. *International Journal for Parasitology* 54, 157–170. https://doi.org/10.1016/j.ijpara.2023.09.003 |
| Hy_asiaticum01 | Jia, N. *et al.* (2020) Large-scale comparative analyses of tick genomes elucidate their genetic diversity and vector capacities. *Cell* 182, 1328–1340. e1313. https://doi.org/10.1016/j.cell.2020.07.023 |

| | |
|---|---|
| I_hexagonus01 | Cerqueira de Araujo, A. *et al.* (2025) Genome sequences of four *Ixodes* species expands understanding of tick evolution. *BMC biology* 23, 17. https://doi.org/10.1186/s12915-025-02121-1 |
| I_inopinatus01 | Baede, V.O. *et al.* (2024) Similarities between *Ixodes ricinus* and *Ixodes inopinatus* genomes and horizontal gene transfer from their endosymbionts. *Current Research in Parasitology & Vector-Borne Diseases* 6, 100229. https://doi.org/10.1016/j.crpvbd.2024.100229 |
| I_inopinatus02 | Baede, V.O. *et al.* (2024) Similarities between *Ixodes ricinus* and *Ixodes inopinatus* genomes and horizontal gene transfer from their endosymbionts. *Current Research in Parasitology & Vector-Borne Diseases* 6, 100229. https://doi.org/10.1016/j.crpvbd.2024.100229 |
| I_inopinatus03 | Baede, V.O. *et al.* (2024) Similarities between *Ixodes ricinus* and *Ixodes inopinatus* genomes and horizontal gene transfer from their endosymbionts. *Current Research in Parasitology & Vector-Borne Diseases* 6, 100229. https://doi.org/10.1016/j.crpvbd.2024.100229 |
| I_pacificus01 | Cerqueira de Araujo, A. *et al.* (2025) Genome sequences of four *Ixodes* species expands understanding of tick evolution. *BMC biology* 23, 17. https://doi.org/10.1186/s12915-025-02121-1 |
| I_persulcatus01 | Jia, N. *et al.* (2020) Large-scale comparative analyses of tick genomes elucidate their genetic diversity and vector capacities. *Cell* 182, 1328–1340. e1313. https://doi.org/10.1016/j.cell.2020.07.023 |
| I_persulcatus02 | Cerqueira de Araujo, A. *et al.* (2025) Genome sequences of four *Ixodes* species expands understanding of tick evolution. *BMC biology* 23, 17. https://doi.org/10.1186/s12915-025-02121-1 |
| I_ricinus01 | Cramaro, W.J. *et al.* (2015) Integration of Ixodes ricinus genome sequencing with transcriptome and proteome annotation of the naïve midgut. *BMC Genomics* 16, 871. https://doi.org/10.1186/s12864-015-1981-7 |
| I_ricinus02 | Cerqueira de Araujo, A. *et al.* (2025) Genome sequences of four *Ixodes* species expands understanding of tick evolution. *BMC biology* 23, 17. https://doi.org/10.1186/s12915-025-02121-1 |
| I_ricinus03 | No paper in preparation (contact Vechtova, P.). |
| I_ricinus04 | Ronai, I. *et al.* (2024) The repetitive genome of the *Ixodes ricinus* tick reveals transposable elements have driven genome evolution in ticks. *bioRxiv*, 2024.2003.2013.584159. https://doi.org/10.1101/2024.03.13.584159 |
| I_ricinus05 | Ronai, I. *et al.* (2024) The repetitive genome of the *Ixodes ricinus* tick reveals transposable elements have driven genome |

| | |
|---|---|
| | evolution in ticks. *bioRxiv*, 2024.2003.2013.584159. https://doi.org/10.1101/2024.03.13.584159 |
| I_ricinus06 | Baede, V.O. *et al.* (2024) Similarities between *Ixodes ricinus* and *Ixodes inopinatus* genomes and horizontal gene transfer from their endosymbionts. *Current Research in Parasitology & Vector-Borne Diseases* 6, 100229. https://doi.org/10.1016/j.crpvbd.2024.100229 |
| I_ricinus07 | Baede, V.O. *et al.* (2024) Similarities between *Ixodes ricinus* and *Ixodes inopinatus* genomes and horizontal gene transfer from their endosymbionts. *Current Research in Parasitology & Vector-Borne Diseases* 6, 100229. https://doi.org/10.1016/j.crpvbd.2024.100229 |
| I_ricinus08 | Baede, V.O. *et al.* (2024) Similarities between *Ixodes ricinus* and *Ixodes inopinatus* genomes and horizontal gene transfer from their endosymbionts. *Current Research in Parasitology & Vector-Borne Diseases* 6, 100229. https://doi.org/10.1016/j.crpvbd.2024.100229 |
| I_scapularis01 | Gulia-Nuss, M. *et al.* (2016) Genomic insights into the *Ixodes scapularis* tick vector of Lyme disease. *Nature Communications* 7. https://doi.org/10.1038/ncomms10507 |
| I_scapularis02 | Miller, J.R. *et al.* (2018) A draft genome sequence for the *Ixodes scapularis* cell line, ISE6. *F1000Research* 7, 297. https://doi.org/10.12688/f1000research.13635.1 |
| I_scapularis03 | De, S. *et al.* (2023) A high-quality *Ixodes scapularis* genome advances tick science. *Nature Genetics* 55, 301–311. https://doi.org/10.1038/s41588-022-01275-w |
| I_scapularis04 | Nuss, A.B. *et al.* (2023) The highly improved genome of *Ixodes scapularis* with X and Y pseudochromosomes. *Life science alliance* 6. https://doi.org/10.26508/lsa.202302109 |
| R_annulatus01 | Guerrero, F.D. *et al.* (2021) Raw pacific biosciences and illumina sequencing reads and assembled genome data for the cattle ticks *Rhipicephalus microplus* and *Rhipicephalus annulatus*. *Data in Brief* 35, 106852. https://doi.org/10.1016/j.dib.2021.106852 |
| R_appendiculatus01 | Meiring, C. *et al.* (2025) Tick genomics through a Nanopore: a low-cost approach for tick genomics. *BMC Genomics* 26, 591. https://doi.org/10.1186/s12864-025-11733-4 |
| R_microplus01 | Guerrero, F.D. *et al.* (2010) Reassociation kinetics-based approach for partial genome sequencing of the cattle tick, *Rhipicephalus (Boophilus) microplus*. *BMC genomics* 11, 374. https://doi.org/10.1186/1471-2164-11-374 |
| R_microplus02 | Barrero, R.A. *et al.* (2017) Gene-enriched draft genome of the cattle tick *Rhipicephalus microplus*: assembly by the hybrid Pacific Biosciences/Illumina approach enabled analysis of the highly repetitive genome. *International journal for parasitology* 47, 569–583. https://doi.org/10.1016/j.ijpara.2017.03.007 |

| | |
|---|---|
| R_microplus03 | Jia, N. *et al.* (2020) Large-scale comparative analyses of tick genomes elucidate their genetic diversity and vector capacities. *Cell* 182, 1328–1340. e1313. https://doi.org/10.1016/j.cell.2020.07.023 |
| R_microplus04 | Guerrero, F.D. *et al.* (2021) Raw pacific biosciences and illumina sequencing reads and assembled genome data for the cattle ticks *Rhipicephalus microplus* and *Rhipicephalus annulatus*. *Data in Brief* 35, 106852. https://doi.org/10.1016/j.dib.2021.106852 |
| R_microplus05 | Tidwell, J.P. *et al.* (2024) Identifying the sex chromosome and sex determination genes in the cattle tick, *Rhipicephalus (Boophilus) microplus*. *G3 Genes\|Genomes\|Genetics* 14. https://doi.org/10.1093/g3journal/jkae234 |
| R_microplus06 | Meiring, C. *et al.* (2025) Tick genomics through a Nanopore: a low-cost approach for tick genomics. *BMC Genomics* 26, 591. https://doi.org/10.1186/s12864-025-11733-4 |
| R_sanguineus01 | Jia, N. *et al.* (2020) Large-scale comparative analyses of tick genomes elucidate their genetic diversity and vector capacities. *Cell* 182, 1328–1340. e1313. https://doi.org/10.1016/j.cell.2020.07.023 |
